# sampbias, a method for quantifying geographic sampling biases in species distribution data

**DOI:** 10.1101/2020.01.13.903757

**Authors:** Alexander Zizka, Alexandre Antonelli, Daniele Silvestro

## Abstract

Geo-referenced species occurrences from public databases have become essential to biodiversity research and conservation. However, geographical biases are widely recognized as a factor limiting the usefulness of such data for understanding species diversity and distribution. In particular, differences in sampling intensity across a landscape due to differences in human accessibility are ubiquitous but may differ in strength among taxonomic groups and datasets. Although several factors have been described to influence human access (such as presence of roads, rivers, airports and cities), quantifying their specific and combined effects on recorded occurrence data remains challenging. Here we present *sampbias*, an algorithm and software for quantifying the effect of accessibility biases in species occurrence datasets. *Sampbias* uses a Bayesian approach to estimate how sampling rates vary as a function of proximity to one or multiple bias factors. The results are comparable among bias factors and datasets. We demonstrate the use of *sampbias* on a dataset of mammal occurrences from the island of Borneo, showing a high biasing effect of cities and a moderate effect of roads and airports. *Sampbias* is implemented as a well-documented, open-access and user-friendly R package that we hope will become a standard tool for anyone working with species occurrences in ecology, evolution, conservation and related fields.

## Background

Publicly available datasets of geo-referenced species occurrences, such as provided by the Global Biodiversity Information Facility (www.gbif.org) have become a fundamental resource in biological sciences, especially in biogeography, conservation, and macroecology. However, these datasets are typically not collected systematically and rarely include information on collection effort. Instead, they are often compiled from a variety of sources (e.g. scientific expeditions, census counts, genetic barcoding studies, and citizen-science observations). Species occurrences are therefore often subject to multiple sampling biases (Meyer et al. 2016).

Sampling biases that may affect the recording of species occurrences (presence, absence and abundance, Isaac and Pocock 2015, Boakes et al. 2010) include the under-sampling of specific taxa (“taxonomic bias”, e.g., birds *vs.* nematodes), specific geographic regions (“geographic bias”, i.e. easily accessible *vs.* remote areas), and specific temporal periods (“temporal bias”, i.e. wet season *vs.* dry season). In particular geographic sampling bias—the fact that sampling effort is spatially biased, rather than equally distributed over the study area—is likely to be widespread in all non-systematically collected datasets of species distributions.

Many aspects can lead to sampling biases, including socio-economic factors (i.e. national research spending, history of scientific research; www.bio-dem.surge.sh, Meyer et al. 2015, Daru et al. 2018), political factors (armed conflict, democratic rights; Rydén et al. 2019), and physical accessibility (i.e. distance to a road or river, terrain conditions, slope; Yang et al. 2014, Botts et al. 2011). Especially physical accessibility by people is omnipresent as a bias factor (e.g. Lin et al. 2015, Kadmon et al. 2004, Engemann et al. 2015), across spatial scales, as the commonly used term “roadside bias” testifies. In practice, this means that most species observations are made in or near cities, along roads, paths, and rivers, and near human settlements. Relatively fewer observations are expected to be available from inaccessible areas in e.g. a tropical rainforest or a mountain top. Since the recording of different taxonomic groups poses different challenges, geographic sampling bias and the effect of accessibility may differ among taxonomic groups (Vale and Jenkins 2012).

The implications of not considering geographic sampling biases in biodiversity research are likely to be substantial (Rocchini et al. 2011, Barbosa et al. 2013, Yang et al. 2013, Kramer-Schadt et al. 2013, Shimadzu and Darnell 2015, Meyer et al. 2016). The presence of geographic sampling biases is broadly recognized (e.g. Kadmon et al. 2004), and approaches exist to account for it in some analyses—such as for species-richness estimates (Engemann et al. 2015) species distribution models (Beck et al. 2014, Varela et al. 2014, Warren et al. 2014, Boria et al. 2014, Fourcade et al. 2014, Fithian et al. 2015, Stolar and Nielsen 2015, Monsarrat et al. 2019), occupancy models (Kery and Royle 2016), and abundance estimates (Shimadzu and Darnell 2015). In contrast, few attempts have been made to explicitly quantify the overall bias (Hijmans et al. 2000, Kadmon et al. 2004) or to discern and quantify different sources of bias (Fithian et al. 2015, Fernández and Nakamura 2015, Ruete 2015). To our knowledge, no tools exist for comparing the strength of bias factors or datasets. We define as *bias factors* any anthropogenic or natural features that facilitate human access and sampling, such as roads, rivers, airports, and cities.

It is unrealistic to expect that accessibility bias in biodiversity data will ever disappear even after more automated observation technologies are developed. It is therefore crucial that researchers realise the intrinsic biases associated with the data they deal with. This is the first step towards estimating to which extent these biases may affect their analyses, results, and conclusions. Any study dealing with species occurrence data should arguably assess the strength of accessibility biases in the underlying data. Such a quantification can also help researchers to target further sampling efforts.

Here, we present *sampbias*, a probabilistic method to quantify accessibility bias in datasets of species occurrences. *Sampbias* is implemented as a user-friendly R-package and uses a Bayesian approach to address three questions:

1. How strong is the accessibility bias in a given dataset?
2. How strong is the effect of different bias factors in causing the overall accessibility bias?
3. How is accessibility bias distributed in space, i.e. which areas are a priority for targeted sampling?

*Sampbias* is implemented in R (R Core Team 2019), based on commonly used packages for data handling (`ggplot`, Wickham 2009, `forcats`, 2019, `tidyr`, Wickham and Henry 2019, `dplyr`, Wickham et al. 2019, `magrittr`, Bache and Wickham 2014, `viridis`, Garnier 2018), handling geographic information and geo-computation (`raster`, Hijmans 2019, `sp`, Pebesma and Bivand 2005, Bivand et al. 2013) and statistical modelling (`stats`, R Core Team 2019). *Sampbias* offers an easy and largely automated means for biodiversity scientists and non-specialists alike to explore bias in species occurrence data, in a way that is comparable across datasets. The results may be used to identify priorities for further collection or digitalization efforts, improve species distribution models (by providing bias surfaces in the analyses), or assess the reliability of scientific results based on publicly available species distribution data.

## Methods and Features

### General concept

Under the assumption that organisms exist across the entire area of interest, we can expect the number of sampled occurrences in a restricted area, such as a single biome, to be distributed uniformly in space (even though, of course, the density of individuals and the species diversity may be heterogeneous). With *sampbias* we assess to which extent variation in sampling rates can be explained by distance from bias factors.

*Sampbias* works at a user-defined spatial scale, and any dataset of multi-species occurrence records can be tested against any geographic gazetteer. Reliability increases with increasing dataset size. Default global gazetteers for airports, cities, rivers and roads are provided with *sampbias*, and user-defined gazetteers can be added easily. Species occurrence data as downloaded from the data portal of GBIF can be directly used as input data for *sampbias*. The output of the package includes measures of the sampling rates across space, which are comparable between different gazetteers (e.g. comparing the biasing effect of roads and rivers), different taxa (e.g. birds *vs.* flowering plants) and different data sets (e.g. specimens *vs.* human observations).

### Distance calculation

*Sampbias* uses gazetteers of the geographic location of bias factors (hereafter indicated with B) to generate a regular grid across the study area (the geographic extent of the dataset). For each grid cell *i*, we then compute a vector *X_i_*(*j*) of minimum distances (straight aerial distance, “as the crow flies”) to each bias factor *j* ∈ *B*. The resolution of the grid defines the precision of the distance estimates, for instance a 1×1 degree raster will yield approximately a 110 km precision at the equator. Due to the assumption of homogeneous sampling and a computational trade-off between the resolution of the regular grid and the extent of the study area (for instance, a 1 second resolution for a global dataset would become computationally prohibitive in most practical cases), *sampbias* is best suited for local or regional datasets at high resolution (c. 100 – 10,000 m).

### Quantifying accessibility bias using a Bayesian framework

We describe the observed number of sampled occurrences *S_i_* within each cell *i* as the result of a Poisson sampling process with rate *λ_i_*. We model the rate *λ*_i_as a function of a parameter *q*, which represents the expected number of occurrences per cell in the absence of biases, i.e. when 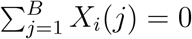. Additionally, we model *λ_i_* to decrease exponentially as a function of distance from bias factors, such that increasing distances will result in a lower sampling rate. For a single bias factor the rates of cell *i* with distance *X_i_* from a bias is:

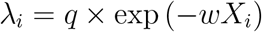

where *w* ∈ ℝ^+^ defines the steepness of the Poisson rate decline, such that *w* ≈ 0 results in a null model of uniform sampling rate *q* across cells. In the presence of multiple bias factors (e.g. roads and rivers), the sampling rate decrease is a function of the cumulative effects of each bias and its distance from the cell:

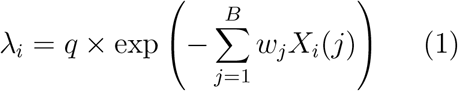

where a vector **w** = [*w*_1_,*…, w_B_*] describes the amount of bias attributed to each specific factor.

To quantify the amount of bias associated with each factor, we jointly estimate the parameters *q* and **w** in a Bayesian framework. We use Markov Chain Monte Carlo (MCMC) to sample these parameters from their posterior distribution:

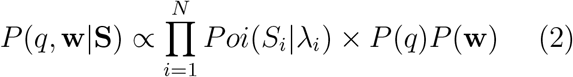

where the likelihood of sampled occurrences *S_i_* within each cell *P oi*(*S_i_*|*λ_i_*) is the probability mass function of a Poisson distribution with rate per cell defined as in Eqn. (1). The likelihood is then multiplied across the *N* cells considered. We used exponential priors on the parameters *q* and **w**, *P* (*q*) ∼ Γ(1, 0.01) and *P* (**w**) ∼ Γ(1, 1), respectively.

We summarize the parameters by computing the mean of the posterior samples and their standard deviation. We interpret the magnitude of the elements in **w** as a function of the importance of the individual biases. We note, however, that this test is not explicitly intended to assess the significance of each bias factor (for which a Bayesian variable selection method could be used), particularly since several bias factors might be correlated (e.g. cities, and airports). Instead, these analyses can be used to quantify the expected amount of bias in the data that can be predicted by single or multiple predictors in order to identify under-sampled and unexplored areas.

We summarize the results by mapping the estimated sampling rates (*λ_i_*) across space. These rates represent the expected number of sampled occurrences for each grid cell and provide a graphical representation of the spatial variation of sampling rates. Provided that the cells are of equal size, the estimated rates will be comparable across data sets, regions, and taxonomic groups. Analysing different regions, biomes, or taxa in separate analyses allows to account for differences in over sampling rates, which are not linked with bias factors. For instance, the unbiased sampling rate *q* is expected to differ between a highly sampled clade like birds and under-sampled groups of invertebrates, but their sampling biases (**w**) might be similar across the two groups.

### Example and Empirical validation

A default *sampbias* analysis can be run with few lines of code in R. The main function `calculate_bias` creates an object of the class `“sampbias”`, for which the package provides a plotting and summary method. Based on a `data.frame` including species identity and geographic coordinates. Additional options exist to provide custom gazetteers, a custom grain size of the analysis, as well as some operators for the calculation of the bias distances. A tutorial on how to use *sampbias* is available with the package and in the electronic supplement of this publication (Appendix S1).

To exemplify the use and output of *sampbias*, we downloaded the occurrence records of all mammals available from the island of Borneo (n = 6,262, GBIF.org 2016), and ran *sampbias* using the default gazetteers as shown in the example code below, to test the biasing effect of the main airports, cities and roads in the dataset. The example dataset is provided with *sampbias*.

We found a strong effect of cities on sampling intensity, a moderate effect of roads and airports and negligible effect of rivers (Fig. 1). All models predict a low number of collection records in the centre of Borneo (Fig. 2), which reflects the original data, and where accessibility means are low (Figure S1 in Appendix S2). The empirical example illustrates the use of *sampbias*, for detailed analyses or a smaller geographic scale, higher resolution gazetteers, including smaller roads and rivers and a higher spatial resolution would be desirable. Results might change with increasing resolution, since roads and rivers might have a stronger effect on higher resolutions (facilitating most the access to their immediate vicinity), whereas cities and airports might have a stronger effect on the larger scale (facilitating access to a larger area).

**Figure 1:**
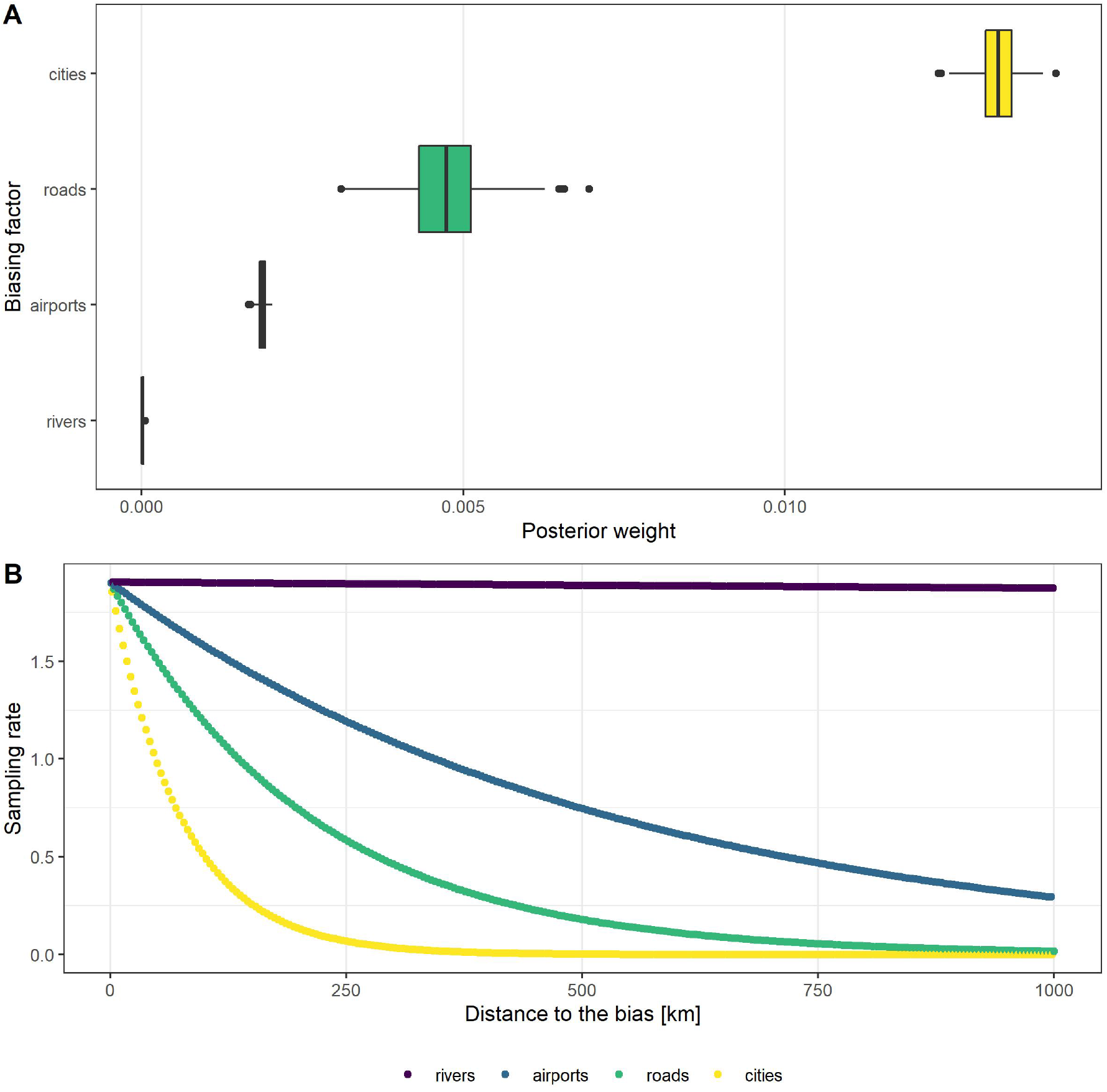
Results of the empirical validation analysis, estimating the accessibility bias in mammal occurrences from Borneo). A) bias weights (*w*) defining the effects of each bias factor, B) sampling rate as function of distance to the closest instance of each bias factor (i.e. expected number of occurrences) given the inferred *sampbias* model. At the study scale of 0.05 degrees (c. 5km) *sampbias* finds the strongest biasing effect for the proximity of cities and roads.

**Figure 2:**
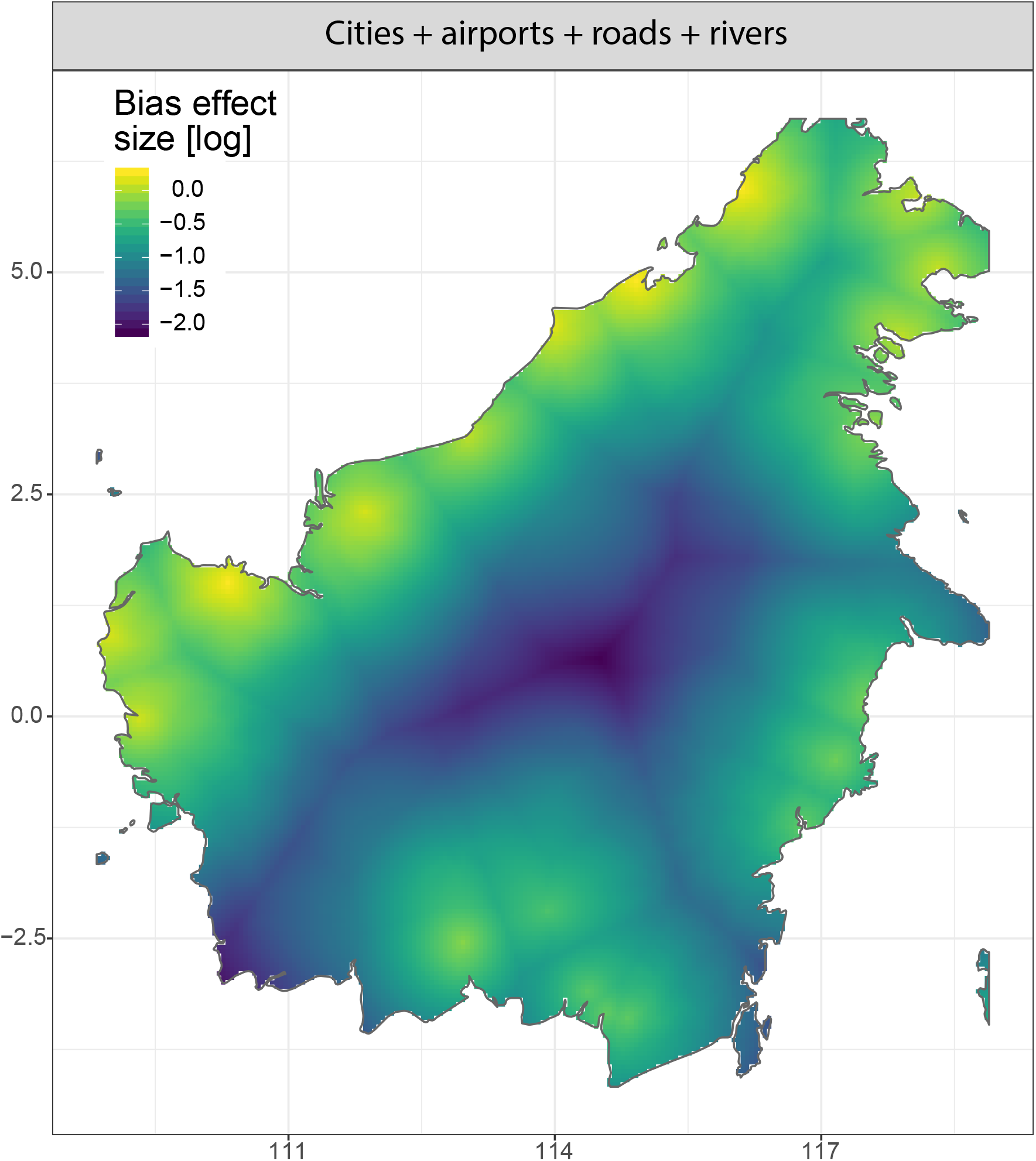
Spatial projection of the estimated sampling rates in an empirical example dataset of mammal occurrences on the Indonesian island of Borneo (downloaded from www.gbif.org. GBIF.org, 2016). The colours show the projection of the sampling rates (i.e. expected number of occurrences per cell) given the inferred extitsampbias model. The highest undersampling is in the centre of the island.

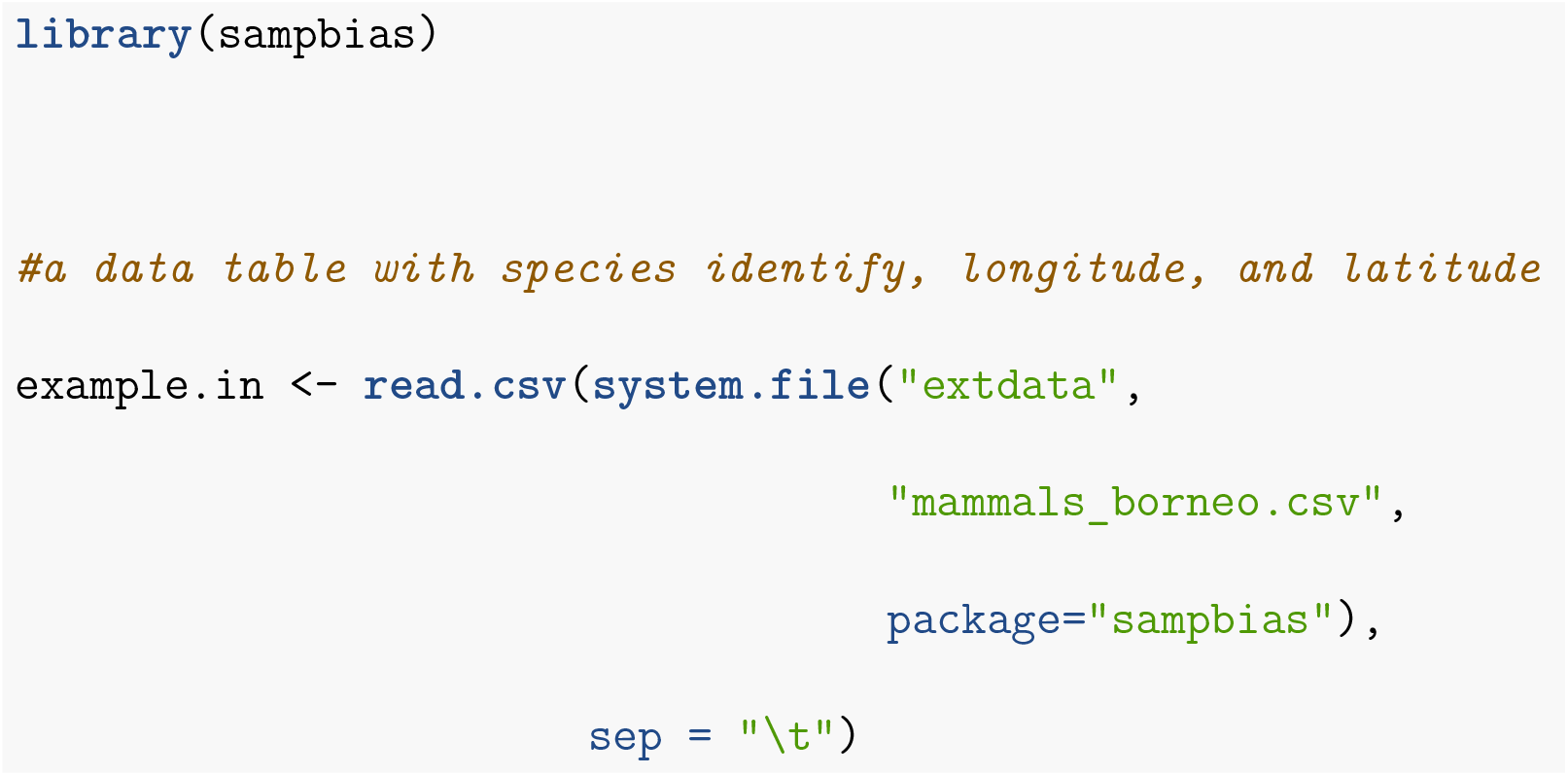
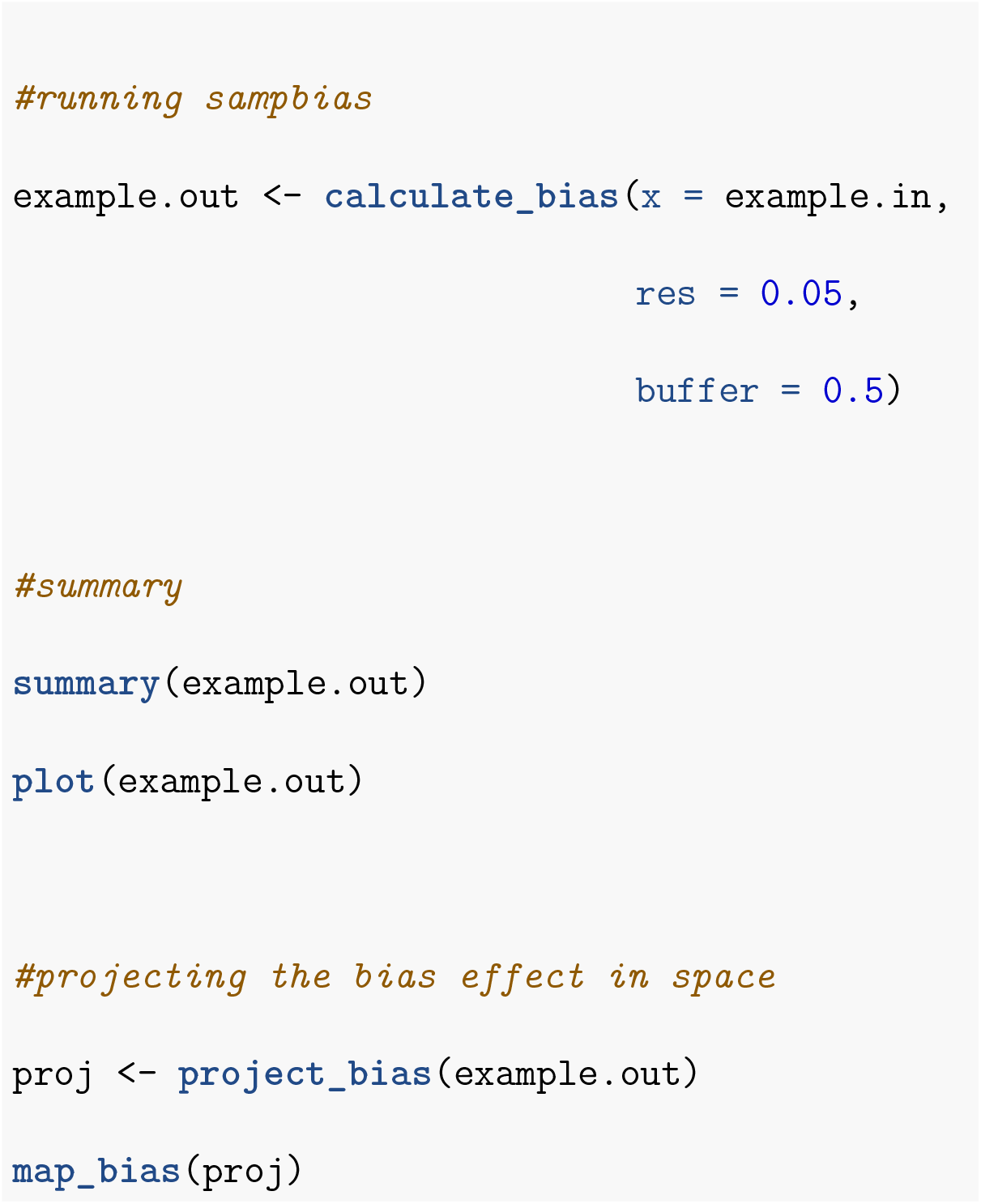

## Supporting information

Appendix S1 - Tutorial running sampbias in R

Appendix S3 - Possible warnings and their solutions

## Data accessibility

*Sampbias* is available under a GNU General Public license v3 from https://github.com/azizka/sampbias, and includes the example dataset as well as a tutorial (Appendix S1) and a summary of possible warnings produced by the package (Appendix S3).

## Acknowledgements

We thank the organizers of the 2016 Ebben Nielsen challenge for inspiring and recognizing this research. We thank all data collectors and contributors to GBIF for their effort. AZ is thankful for funding by iDiv via the German Research Foundation (DFG FZT 118), specifically through sDiv, the Synthesis Centre of iDiv. AA is supported by grants from the Swedish Research Council, the Knut and Alice Wallenberg Foundation, the Swedish Foundation for Strategic Research and the Royal Botanic Gardens, Kew. DS received funding from the Swedish Research Council (2015-04748) and from the Swiss National Science Foundation (PCEFP3_187012)

## Author contributions

All authors conceived this study, AZ and DS developed the statistical algorithm and wrote the R-package, AZ and DS wrote the manuscript with contributions from AA.

## Supplementary material

Appendix S1 - Tutorial running *sampbias* in R

Appendix S2 - Supplementary Figure S1

Appendix S3 - Possible warnings and their solutions

