## Appendix S1 - Tutorial running sampbias in R for "sampbias, a method for quantifying geographic sampling biases in species distribution data"

### Using the sampbias R package

#Installing sampbias To install the *sampbias* R-package, you can either download the latest build version from gitHub use the `devtools` package:

```
require(devtools)
install_github(repo = "azizka/sampbias")
library(sampbias)
library(maptools)
```

#### Input data

*Sampbias* calculates spatial bias in a species occurrence data set based on two input files:

1. A table of Species occurrence records
2. A set of geographical gazetteers

The example files for this tutorial are provided in the example_data folder and should be copied to the R working directory, before starting. You can find descriptions and help for all functions by typing `?Functionname`, for instance `?calculate_bias`.

##### Species occurrence records

Species occurrences can be provided to *sampbias* as an R `data.frame`, which can be loaded from any table format .txt file. The minimum input file needed to run *sampbias* is a table with species occurrences including three columns named "species", "decimallongitude", "decimallatitude". *Sampbias* can also directly work with data downloaded from the Global Biodiversity Information Facility data portal (GBIF). We will use the records of mammals from Borneo provided in the sampbias folder (GBIF, 2016) as example file throughout the tutorial. The file is also included as example data set in the package.

Text files can easily be loaded into R as data.frame using the `read.csv()` function.

```
#loading a text file to R
occ <-read.csv(system.file("extdata",
                           "mammals_borneo.csv",
                           package="sampbias"),
               sep = "\t")
```

**Note**: *Sampbias* evaluates the bias by comparing to a random sampling, meaning that the tool is not designed for single species data sets, as distribution of the records might then reflect ecological preferences, but rather for multi-species data sets. In general, the more species and the more records, the more reliable the results will be.

##### Geographic gazetteers

*Sampbias* evaluates the distribution of the sampled species occurrences in relation to geographic features that might bias sampling effort. These are generally related to accessibility or means of travel. *Sampbias* includes a set of default gazetteers for cities, airports, roads and rivers (Natural Earth, 2016), which can give fair estimates of bias for large and medium scale analyses. These are used by default, if no other gazetteers are provided by the user. However, these defaults include major features only and if available high resolution user-provided gazetteers are preferable.

Any number of gazetteers can be provided to *sampbias* via the `gaz` argument as a list of `SpatialPointsDataFrame` and `SpatialLinesDataFrame` objects. These objects can easily be loaded into R from standard shape files using the `maptools` package:

```
cit <- readShapeSpatial(system.file("extdata",
                                    "Borneo_major_cities.shp",
                                    package="sampbias"))
roa <- readShapeSpatial(system.file("extdata",
                                    "Borneo_major_roads.shp",
                                    package="sampbias"))

gazetteers <- list(cities = cit,
                   roads = roa)
```

See here and `?SpatialPointsDataFrame` or `?SpatialLinesDataFrame` on how to create a SpatialObjects from tables of coordinates.

### Running an analysis

A sampbias analyses is run in one line of code via the `calculate_bias` function:

```
bias.out <- calculate_bias(x = occ, gaz = gazetteers)
```

In addition to the input from above, `calculate_bias` offers a set of options to customize analyses, of which the most important ones are shown in Table 1. See `?calculate_bias` for a detailed description of all options.

| Option | Description |
|---|---|
| res | the raster resolution for the distance calculation to the geographic features and the data visualization in decimal degrees. The default is to one degree, but higher resolution will be desirable for most analyses. Res together with the extent of the input data determine computation time and memory requirements. |
| convexhull | logical. If TRUE, the empirical distribution (and the output maps) is restricted to cells within a convex hull polygon around `x`. If FALSE a rectangle around `x` is used. Default = FALSE. |
| terrestrial | logical. If TRUE, the empirical distribution (and the output maps) are restricted to terrestrial areas. Default = TRUE. |

The default MCMC runs for 100,000 generations with a 20% burn-in, which has proven sufficient for most analyses. We suggest to verify that the effective sample size of the posterior estimates is large enough (e.g. > 200) using the `ESS` function of the `LaplacesDemon` package (`LaplacesDemon::ESS(bias.out$bias_estimate)`).

### Output

The output of `calculate_bias` is a list of different R objects.

| Object | Description |
|---|---|
| summa | A list of summary statistics for the sampbias analyses, including the total number of occurrence points in x, the total number of species in x, the extent of the output rasters as well as the settings for res, binsize, and convexhull used in the analyses. |
| occurrences | a raster indicating occurrence records per grid cell, with resolution res. |

The package includes summary and plot methods for an easy exploration of the results:

```
summary(bias.out)
plot(bias.out)
```

The plot method generates a boxplot of the posterior estimates of the weights for each biasing factor.

As the last step, it is possible to project the bias effects into space and amp them, to identify areas with particular high bias, for instance, to design future field campaigns.

```
proj <- project_bias(bias.out)
map_bias(proj)
```

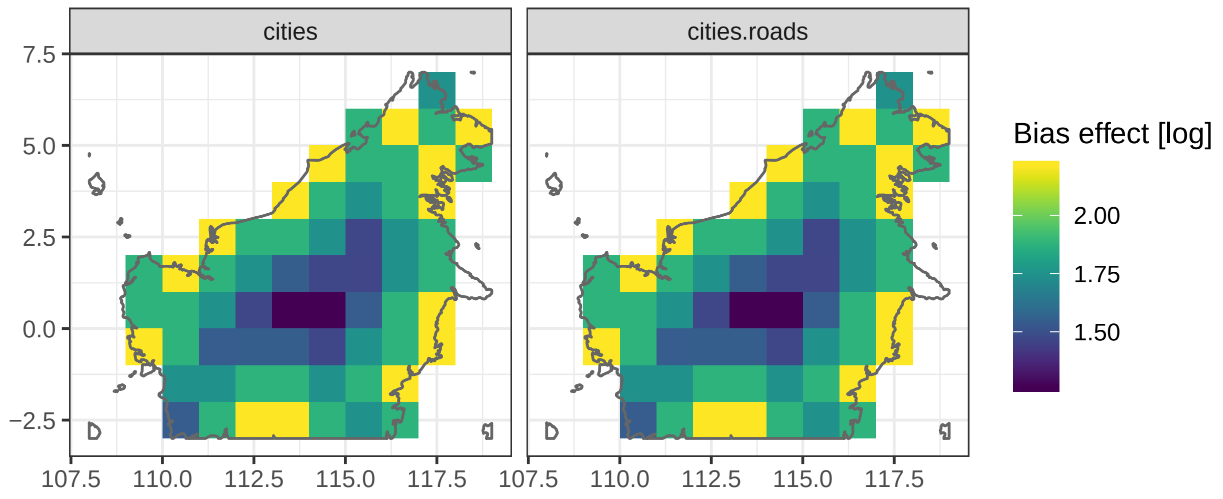
