## Appendix S2 - Supplementary Figure S1 for "sampbias, a method for quantifying geographic sampling biases in species distribution data"

### Appendix S1 - Supplementary Figure

Sampbias, a method to evaluate geographic sampling bias in species distribution data

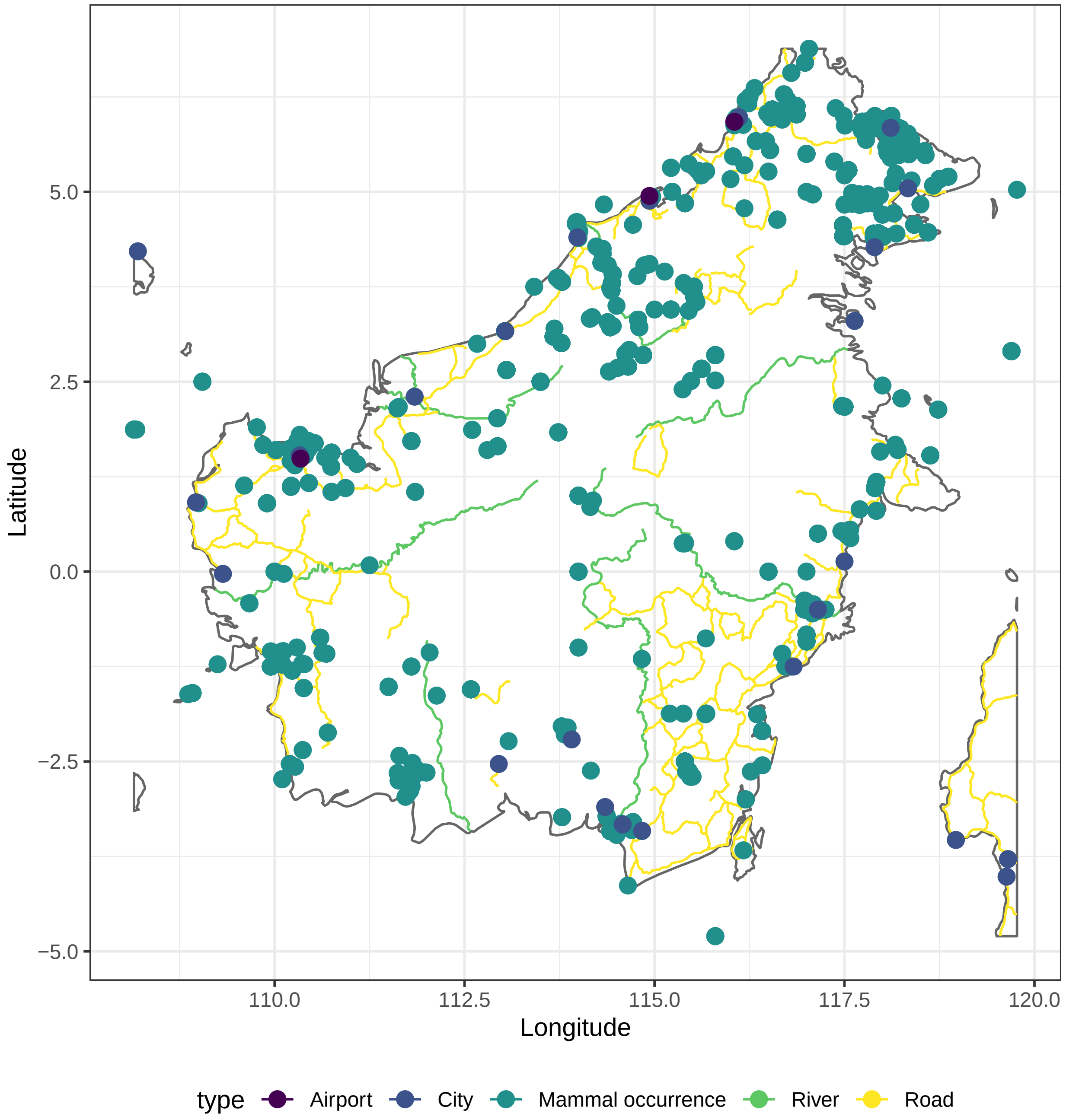


Figure S1: The example dataset of mammal occurrences from the island of Borneo, as downloaded from www.gbif.org (n = 6,262), and the geographic gazetters of main cities, roads, rivers and airports used for the sampbias analysis.
