## Appendix S3 - Possible warnings and their solutions for "sampbias, a method for quantifying geographic sampling biases in species distribution data"

### Possible warnings of sampbias and solutions

Sampbias, might return warning messages depending on input data and settings. All should be fixable, please report additional bugs [here] (https://github.com/azizka/sampbias/issues).

1. *"'gaz' not found, using standard gazetteers"*

    Occurs if the `gaz` option is not specified when calling `SamplingBias`, indicating a fall back to the default gazetteers distributed with the package. This is generally not a problem, and might even be desired. Note however, that the default gazetteers o are of global scale and thus, relatively coarse. So, especially for small scale analyses, more detailed regional to local gazetteers might be desirable. Default gazetteers include airports, cities, rivers and roads. Check here for more information on how to use custom gazetteers.

2. *"Low resolution, increase resolution or switch to absolute distance calculation"*

    Occurs if the number of bins for approximating the statistical distribution is lower than 100. The number of bins is dependent on res, so try to increase res. If an increase of res is not possible (e.g. due to computation time) try to adapt the number of bins directly via `binsize`. However, this might cause other problems and should be avoided. The minimum number of bins necessary for robust results is data set dependent and the threshold of 100 is currently arbitrary. Thus the warning does not necessarily indicated problems with the analysis. A best practice to evaluate if the low resolution is problematic is to switch on the `extraplot` option of `SamplingBias` and visually inspect the statistical distributions which should be as smooth as possible.

3. *"huge raster size"*

    Occurs if the raster of the study area exceeds 1 M grid cells, due to the combination of study area extent and the chosen `res`. There is no theoretical limit to the number of grid cells, but calculations might get slow and hit memory limits. Try do either reduce the extent of the study area or decrease resolution.

4. *"Adapting buffer precision to resolution. Buffer set to X"* or " 'Buffer' is not a multiple of res. Set 'buffer' to a multiple of res"

    Occurs if the `buffer` is not a multiple of `res` which might cause undesired raster shifts. Sampbias automatically changes the buffer to be a multiple of res. To avoid this warning always set the buffer to any desired multiple of res. Usually can be safely ignored.

5. *"Evening buffer. Buffer set to X"*

    Occurs if the `buffer` is set to an odd number. The buffer has to be even, as it added to all sides of the extent of X. To avoid this warnings set `buffer` to an even number. Usually can be safely ignored.

6. *"No reference found for 'X'. Increase buffer."*

    Occurs if for any gazetteer no element is found with the study area, e.g. no road in the entire area. This mostly can happen in small scale studies of remote areas using coarse gazetteers. Increase `buffer` to include the closest element of the gazetteer, or exclude the respective gazetteer from the study. Problematic gazetteers will be ignored for the analyses.

7. *"No references found within study boundaries. Increase buffer. Falling back to species and occurrence raster"*

    Occurs if no elements for any gazetteer can be found in the study area. Increase the buffer or provide a more suitable set of gazetteers. Only occurrence and species raster are produced as output.

8. *"Gazetteers have NA values in first column, the corresponding entries will be ignored"*

Occurs if there are any NA entries in the `data` slot of the `SpatialPointsDataFrame` or `SpatialLinesDataFrame` objects used as gazetteers. Check the gazetteers and replace NA values with any value.
